# Suppressed sensory response to predictable object stimuli throughout the ventral visual stream

**DOI:** 10.1101/228890

**Authors:** David Richter, Matthias Ekman, Floris P. de Lange

## Abstract

Prediction plays a crucial role in perception, as prominently suggested by predictive coding theories. However, the exact form and mechanism of predictive modulations of sensory processing remain unclear, with some studies reporting a downregulation of the sensory response for predictable input, while others observed an enhanced response. In a similar vein, downregulation of the sensory response for predictable input has been linked to either sharpening or dampening of the sensory representation, which are opposite in nature. In the present study we set out to investigate the neural consequences of perceptual expectation of object stimuli throughout the visual hierarchy, using fMRI in human volunteers. Participants (n=24) were exposed to pairs of sequentially presented object images in a statistical learning paradigm, in which the first object predicted the identity of the second object. Image transitions were not task relevant; thus all learning of statistical regularities was incidental. We found strong suppression of neural responses to expected compared to unexpected stimuli throughout the ventral visual stream, including primary visual cortex (V1), lateral occipital complex (LOC), and anterior ventral visual areas. Expectation suppression in LOC, but not V1, scaled positively with image preference, lending support to the dampening account of expectation suppression in object perception.

**Significance Statement:** Statistical regularities permeate our world and help us to perceive and understand our surroundings. It has been suggested that the brain fundamentally relies on predictions and constructs models of the world in order to make sense of sensory information. Previous research on the neural basis of prediction has documented expectation suppression, i.e. suppressed responses to expected compared to unexpected stimuli. In the present study we queried the presence and characteristics of expectation suppression throughout the ventral visual stream. We demonstrate robust expectation suppression in the entire ventral visual pathway, and underlying this suppression a dampening of the sensory representation in object-selective visual cortex, but not in primary visual cortex. Taken together, our results provide novel evidence in support of theories conceptualizing perception as an active inference process, which selectively dampens cortical representations of predictable objects. This dampening may support our ability to automatically filter out irrelevant, predictable objects.

## Introduction

Our environment is structured by statistical regularities. Making use of such regularities by anticipating upcoming stimuli is of great evolutionary value, as it enables the agent to predict future states of the world and prepare adequate responses, which in turn can be executed faster or more accurately (Bertels et al., 2012; Hunt and Aslin, 2001; Kim et al., 2009). Our brains are exquisitely sensitive to these statistical regularities (Schapiro et al., 2012; Schapiro et al., 2014; Turk-Browne et al., 2009; Turk-Browne et al., 2010). In fact, it has been suggested that a core operational principle of the brain is prediction (Bubic et al., 2010) and prediction error minimization (Friston, 2005). Statistical learning is an automatic learning process by which statistical regularities are extracted from the environment (Turk-Browne et al., 2010), without explicit awareness or effort by the observer (Brady and Oliva, 2008; Fiser and Aslin, 2002), even under concurrent cognitive load (Garrido et al., 2016). These statistical regularities can be used to form predictions about upcoming input, with effects of statistical learning being evident even 24 hours after exposure (Kim et al., 2009).

The neural consequences of perceptual predictions have been investigated extensively, but conflicting results have emerged. For example, Turk-Browne et al. (2009) reported larger neural responses to predictable than random sequences of stimuli in human object-selective lateral occipital complex (LOC). However, contrary to this notion, neurons in monkey inferotemporal cortex (IT), the putative homologue of human LOC (Denys et al., 2004), showed reduced responses to expected compared to unexpected object stimuli (Meyer and Olson, 2011; Kaposvari et al., 2016). This is in line with findings in human primary visual cortex (V1), which revealed that visual gratings of an expected orientation elicit a suppressed neural response compared to gratings of an unexpected orientation (Kok et al., 2012a; St. John-Saaltink et al., 2015). Even though there is superficial agreement between these studies, the exact form of expectation suppression appeared to be opposite. Kok et al. (2012a) observed the strongest suppression in voxels that were tuned away from the expected stimulus, resulting in a sparse, sharpened population code. Electrophysiological studies in macaques on the other hand have reported a positive scaling of expectation suppression with image preference (Meyer and Olson, 2011), suggesting that sensory representations are dampened for expected stimuli (Kumar et al., 2017).

In sum, several discrepancies remain concerning the neural basis of perceptual expectation, which may be related to differences in species (macaque vs. human), cortical hierarchy (early vs. late) and measurement technique (spike rates vs. fMRI BOLD). In the current study, we set out to examine the existence and characteristics of expectation suppression throughout the visual hierarchy, using a paradigm that closely matches previous literature on object prediction in macaque monkeys (Meyer and Olson, 2011; Ramachandran et al., 2016). This allows us to better compare and generalize between species and methods, while measuring neural activity throughout the entire human brain. First we exposed participants to pairs of sequentially presented object images in a statistical learning paradigm. Next, we recorded neural responses, using whole-brain fMRI, to expected and unexpected object image pairs. By contrasting responses to expected and unexpected pairs we probed whether a suppression of expected object stimuli is evident throughout the ventral visual stream, and in particular in object-selective cortex. Moreover, by investigating expectation suppression as a function of image preference we contrasted sharpening against dampening (scaling) accounts of expectation suppression.

In brief, our results show that expectation suppression is ubiquitous throughout the human ventral visual stream, including object-selective LOC. Furthermore, we found that expectation suppression positively scales with object image preference within object-selective LOC, but not V1. This suggests that object predictions selectively dampen sensory representations in object-selective regions.

## Materials and Methods

### Participants

Twenty-four healthy, right-handed participants (17 female, aged 23.3 ± 2.4 years, mean ± SD) were recruited from the Radboud research participation system. The sample size was based on an a priori power calculation, computing the required sample size to achieve a power of 0.8 to detect an effect size of Cohen’s d = 0.6, at alpha = 0.05, for a two-tailed within subjects t-test. Participants were prescreened for MRI compatibility, had no history of epilepsy or cardiac problems, and normal or corrected-to-normal vision. Written informed consent was obtained before participation. The study followed institutional guidelines of the local ethics committee (CMO region Arnhem-Nijmegen, The Netherlands). Participants were compensated with 42 euro for study participation. Data from one subject was excluded due to excessive tiredness and poor fixation behavior. One additional subject was excluded from all ROI based analyses, since no reliable object-selective LOC mask could be established due to subpar fixation behavior during the functional localizer.

### Experimental Design and Statistical Analysis

#### Stimuli and experimental paradigm

##### Main task

Participants were exposed to two object images in quick succession. Each image was presented for 500 ms without interstimulus interval, and an intertrial interval of 1500-2500 ms during behavioral training and 4110-6300 ms during fMRI scanning (see Figure 1A for a single trial). A fixation bullseye (0.5° visual angle in size) was presented throughout the run. For each participant 16 object images were randomly selected from a pool of 80 stimuli (also see: *Stimuli*). Eight images were assigned as leading images, i.e. appearing first on trials, while the other eight images served as trailing images, occurring second. Image pairs and the transitional probabilities between them were determined by the transitional probability matrix depicted in Figure 1B, based on the transition matrix used by Ramachandran et al. (2016). The expectation manipulation consisted of a repeated pairing of images in which the leading image predicted the identity of the trailing image, thus over time making the trailing image expected given the leading image. Importantly, the transitional probabilities governing the associations between images were task irrelevant, since participants were instructed to respond, by button press, to any upside-down versions of the images, the occurrence of which was not related to the transitional probability manipulation and could not be predicted. Upside-down images (target trials) occurred on ^~^9% of trials. Participants were not informed about the presence of any statistical regularities and instructed to maintain fixation on the central fixation bulls-eye. Trial order was fully randomized.

**Figure 1.**
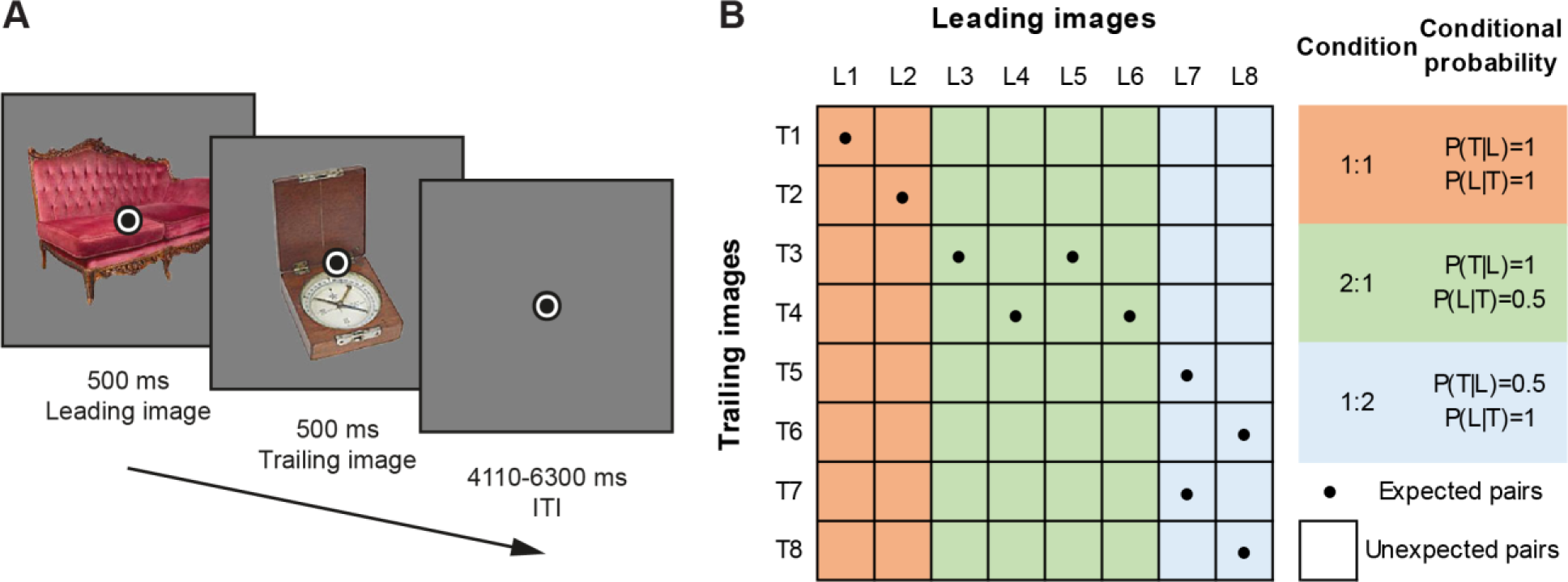
Paradigm overview. (**A**) Depicts a single trial, with two example images and superimposed fixation bullseye. Leading images and trailing images were presented for 500 ms each, without interstimulus interval, followed by an intertrial interval of 4110-6300 ms (fMRI session; 1500-2500 ms during behavioral training). Participants responded to upside-down images by button press; the image at either position (leading or trailing) could be upside-down. (**B**) Shows the utilized image transition matrix determining image pairs. Eight leading images (L1 - L8) and eight trailing images (T1-T8) were used for each participant. Conditional probability conditions are highlighted and their respective conditional probabilities during training are listed on the right; condition 1:1 (orange), 2:1 condition (green), 1:2 condition (blue). Cells with dots indicate expected image pairs, while empty cells denote unexpected pairs.

During behavioral training only expected image pairs were presented on a total of 1792 trials, split into 8 blocks with short breaks in between blocks. Thus, during this session the occurrence of image L1 was perfectly predictive of image T1 (i.e. *P*(T1|L1) = 1; see Figure 1B). Apart from these trials, which constituted the 1:1 conditional probability condition, there were also trials with a 2:1 and 1:2 image pairing. In the 2:1 conditional probability condition the leading image was perfectly predictive of the trailing image (e.g. *P*(T3|L3) = 1), but two different leading images predicted the same trailing image, thereby reducing the conditional probability of the *leading* image given a particular trailing image (i.e. *P*(L3|T3) = 0.5). Lastly, the 1:2 condition consisted of a reduced predictive probability of the trailing image given the leading image, as such image L7 for instance was equally predictive of images T5 and T7 (i.e. *P*(T5|L7) = 0.5 and *P*(T7|L7) = 0.5).

On the next day participants performed one additional behavioral training block, consisting of 224 trials, and another 48 practice trials in the MRI during acquisition of the anatomical image. The task during the subsequent fMRI experiment was identical to the training session, except that also unexpected image pairs occurred. Nonetheless, the expected trailing image was still most likely to follow a given leading image, namely on 56.25% of trials compared to 6.25% for each unexpected trailing image (1:1 condition). Since intertrial intervals were longer in the fMRI session, and responses to upside-down images therefore occurred at a lower rate, potentially reducing participants’ vigilance, the percentage of upside-down images was increased to ^~^11% of trials. As during the behavioral training session, in the main fMRI task participants were not informed about the presence of transitional probabilities, and there was no correlation between the image transitions and the occurrence of upside-down images. In total the MRI main task consisted of 512 trials, split into four equal runs, with an additional three resting blocks (each 12 sec) per run. Feedback on behavioral performance (percent correct and mean response time) was provided after each run. To ensure adequate fixation on the fixation bullseye, an infrared eye tracker (SensoMotoric Instruments, Berlin, Germany) was used to record and monitor eye positions.

##### Functional localizer

The main task was followed by a functional localizer, which was used for a functional definition of object-selective LOC for each participant, and to determine image preference for each voxel within visual cortex in an expectation neutral context. Finally, localizer data served as independent training data for the multi-voxel pattern analysis (see: *Data analysis, Multi-voxel pattern analysis*). In a block design each object image was presented four times, each time flashing at 2 Hz (300 ms on, 200 ms off) for 11 sec. Additionally, a globally phase-scrambled version of each image (Coggan et al., 2016) was shown twice, also flashing at 2 Hz for 11 sec. The order of objects images and scrambles was randomized. Participants were instructed to fixate the bullseye and respond by button press whenever the fixation bullseye dimmed in brightness.

##### Questionnaire

Following the fMRI session, participants filled in a brief questionnaire probing their explicit knowledge of the image transitions. Knowledge of each of the eight image pairs was tested by presenting participants with one leading image at a time, instructing them to select the most likely trailing image.

##### Categorization task

During the categorization task participants were instructed to indicate, by button press, whether the trailing image would fit into a shoebox (yes/no decision); similar to Dobbins et al. (2004), and Horner and Henson (2008). This task was aimed at assessing any implicit reaction time or accuracy benefits due to incidental learning, since in principle the statistical regularities could be used to predict the correct response before the trailing image appeared. For each participant it was ensured that half of the trailing images in each conditional probability condition (1:1, 1:2, 2:1) fit into a shoebox, while the other half did not fit. A brief practice block was used to make sure that participants correctly classified the object images and understood the task. Participants were not informed about the intention behind this task, nor were they instructed to make use of the statistical regularities, in order to avoid influencing their behavior. A full debriefing took place after the categorization task.

##### Stimuli

Object stimuli were taken from Brady et al. (2008), and consisted of a large collection of diverse full-color photographs of objects. Of this full set of images, a subset of 80 images was selected; 40 objects fitting into a shoebox, and 40 objects not fitting into a shoebox. Images spanned approximately 5° × 5° visual angle and were presented in full-color on a mid-grey background. During training stimuli were displayed on a LCD screen and back-projected during MRI scanning (EIKI LC-XL100 projector; 1024 × 768 pixel resolution, 60 Hz refresh rate), visible using an adjustable mirror. Since images were drawn at random per participant, each image could occur in any condition or position, thereby eliminating potential effects induced by individual image features.

#### fMRI data acquisition

Functional and anatomical images were collected on a 3T Skyra MRI system (Siemens, Erlangen, Germany), using a 32-channel headcoil. Functional images were acquired using a whole-brain T2*-weighted multiband-8 sequence (time repetition [TR] / time echo [TE] = 730/37.8 ms, 64 slices, voxel size 2.4 mm isotropic, 50° flip angle, A/P phase encoding direction). Anatomical images were acquired with a T1-weighted magnetization prepared rapid gradient echo sequence (MP-RAGE; GRAPPA acceleration factor = 2, TR/TE = 2300/3.03 ms, voxel size 1 mm isotropic, 8° flip angle).

#### Data analysis

##### Behavioral data analysis

Behavioral data from the categorization task was analyzed in terms of reaction time (RT) and accuracy. All RTs exceeding 3 SD above mean and below 200 ms were excluded as outliers (2.0% of trials). Since unexpected trailing image trials during the categorization task may require a change in the response, any differences in RT and accuracy between the expected and unexpected conditions may reflect a combination of surprise and response adjustment, thereby inflating possible RT and accuracy differences. Therefore, only unexpected trials requiring the same response as the expected image were analyzed, yielding an unbiased comparison of the effect of expectation. RTs for expected and unexpected trailing image trials were averaged separately per participant and subjected to a paired t-test. The error rate was also calculated separately for expected and unexpected trailing image trials per subject and analyzed with a paired t-test. Additionally, the effect size of both differences was calculated in terms of Cohen’s d_z_ (Lakens, 2013). All standard errors of the mean presented here were calculated as the within-subject normalized standard error (Cousineau, 2005) with Morey’s (2008) bias correction.

##### fMRI data preprocessing

fMRI data preprocessing was performed using FSL 5.0.9 (FMRIB Software Library; Oxford, UK; www.fmrib.ox.ac.uk/fsl; Smith et al., 2004). The preprocessing pipeline included brain extraction (BET), motion correction (MCFLIRT), temporal high-pass filtering (128 s), and spatial smoothing for univariate analyses (Gaussian kernel with full-width at half-maximum of 5 mm). No smoothing was applied for multivariate analyses, nor for the voxel-wise image preference analysis. Functional images were registered to the anatomical image using FLIRT (BBR) and to the MNI152 T1 2mm template brain (linear registration with 12 degrees of freedom). The first eight volumes of each run were discarded to allow for signal stabilization.

##### Univariate data analysis

To investigate expectation suppression across the ventral visual stream, voxel-wise general linear models (GLM) were fit to each subject’s run data in an event-related approach using FSL FEAT. Separate regressors for expected and unexpected image pairs were modeled within the GLM. All trials were modeled with one second duration (corresponding to the duration of the leading and trailing image combined) and convolved with a double gamma haemodynamic response function. Additional nuisance regressors were added, including one for target trials (upside-down images), instruction and performance summary screens, first-order temporal derivatives for all modeled event types, and 24 motion regressors (six motion parameters, the derivatives of these motion parameters, the squares of the motion parameters, and the squares of the derivatives; comprising FSL’s standard + extended set of motion parameters). The contrast of interest for the whole-brain analysis compared the average BOLD activity during unexpected minus expected trials, i.e. expectation suppression. Data was combined across runs using FSL’s fixed effect analysis. For the across participants whole-brain analysis, FSL’s mixed effect model FLAME 1 was utilized. Multiple comparison correction was performed using Gaussian random-field based cluster thresholding, as implemented in FSL, using a cluster-forming threshold of z > 3.29 (i.e. *p* < 0.001, two-sided) and a cluster significance threshold of *p* < 0.05. An identical analysis was performed to assess the influence of the different conditional probability conditions (see: *Main task),* except that the expected and unexpected event regressors were split into their respective conditional probability conditions (1:1, 1:2, 2:1), thus resulting in a GLM with six regressors of interest.

#### Planned region of interest analyses

Within each ROI (V1 and LOC; see: *Region of interest definition*), the parameter estimates for the expected and unexpected image pairs were extracted separately from the whole-brain maps. Per subject the mean parameter estimate within the ROIs was calculated and divided by 100 to yield an approximation of mean percent signal change compared to baseline (Mumford, 2007). These mean parameter estimates were in turn subjected to a paired t-test and the effect size of the difference calculated (Cohen’s d_z_). For the conditional probability manipulation, a similar ROI analysis was performed, except that the resulting mean parameter estimates were subjected to a 3x2 repeated measures ANOVA with conditional probability condition (1:1, 2:1, 1:2) and expectation (expected, unexpected) as factors.

### Multi-voxel pattern analysis

Multi-voxel pattern analysis (MVPA) was performed per subject on mean parameter estimate maps per trailing image. These maps were obtained by fitting voxel-wise GLMs per trial for each subject, following the ‘least squares separate’ approach outlined in Mumford et al. (2012). In brief, a GLM is fit for each trial, with only that trial as regressor of interest and the remaining trials as one regressor of no interest. This was done for the functional localizer and main task data. The resulting parameter estimate maps of the functional localizer were used as training data for a multi-class SVM (classes being the eight trailing images), as implemented in Scikit-learn (SVC; Pedregosa et al., 2011).

Decoding performance was tested per subject on the mean parameter estimate maps from the main task data for each trailing image, split into expected and unexpected image pairs. The choice to decode mean parameter estimate maps, instead of single trial estimates, was made after observing that image decoding performance when decoding individual trials was close to chance, indicating a lack of sensitivity to detect potential differences between expected and unexpected image pairs. This decision was based on an independent MVPA collapsed over expected and unexpected image pairs, without inspection of the contrast of interest. Expected image pair trials are by definition more frequent, which may in turn yield a more accurate mean parameter estimate. Thus, stratification by random sampling was used to balance the number of expected and unexpected image pairs per trailing image, thereby removing potential bias. In short, for each iteration (n = 1,000) a subset of expected trials was randomly sampled to match the number of unexpected occurrences of that trailing image. Finally, decoding performance was analyzed in terms of mean decoding accuracy. To this end, the class with the highest probability for each test item was chosen as the predicted class and the proportion of correct predictions calculated. Mean decoding performances for expected and unexpected image pairs were subjected to a two-sided, one sample t-test against chance decoding performance (chance level = 12.5%). If decoding was above chance for the expected and unexpected image pairs, decoding performances between expected and unexpected pairs were compared by means of a paired t-test and the effect size was calculated.

### Image preference analysis

For the voxel-wise image preference analysis the single trial GLM parameter estimate maps, as outlined in the *MVPA* section above, were utilized. Within each participant the parameter estimate maps of the functional localizer were averaged for each trailing image, thus yielding an average activation map induced by each trailing image in an expectation free, neutral context. The same was done for the main task data, but for expected and unexpected occurrence of each trailing image separately. Then, for each voxel, trailing images were ranked according to the response they elicited during the functional localizer. These rankings were applied to the main task data, resulting in a vector per voxel, consisting of the mean activation (parameter estimate) elicited by the trailing images during the main task, ranked from the least to most preferred image based on the context neutral, independent functional localizer data. This was done separately for expected and unexpected occurrence of each trailing image. Within each ROI the mean parameter estimates of expected and unexpected image pairs per preference rank was calculated. For each ROI linear regressions were fit to the ranked parameter estimates, one for expected and one for unexpected pairs. A positive regression slope would thus indicate that the ranking from the functional localizer generalized to the main task, which was considered a prerequisite for any further analysis. This was tested by subjecting the slope parameters across subjects to a one sample t-test, comparing the obtained slopes against zero. If this requirement was met for expected and unexpected slopes, the difference between slope parameters was compared by a paired t-test. If the amount of expectation suppression (i.e. unexpected minus expected) indeed scales with image preference (i.e. dampening), then we should find the slope parameter for the unexpected condition regression line to be significantly larger than for the expected condition. The opposite prediction, a larger slope parameter for the expected condition, is made by the sharpening account. For this comparison the effect size was also calculated in terms of Cohen’s d_z_.

### Region of interest definition

The two a-priori regions of interest, object-selective LOC and V1, were defined per subject based on data that was independent from the main task. In order to obtain object-selective LOC, GLMs were fit to the functional localizer data of each subject, modelling object image and scrambled image events separately with a duration corresponding to their display duration. First-order temporal derivatives, instruction and performance summary screens, as well as motion regressors were added as nuisance regressors. The contrast, object images minus scrambles, thresholded at z > 5 (uncorrected; i.e. p < 1e-5), was utilized to select regions per subject selectively more activated by intact object images compared to scrambles (Kourtzi and Kanwisher, 2001; Haushofer et al., 2008). The threshold was lowered on a per subject basis, if the LOC mask contained less than 300 voxels in native volume space. The individual functional masks were constrained to anatomical LOC using an anatomical LOC mask obtained from the Harvard-Oxford cortical atlas, as distributed with FSL. Finally, a decoding analysis of object images (also see: *Multi-voxel pattern analysis*) was performed using a searchlight approach (6 mm radius) on the functional localizer data, using a k-fold cross-validation scheme with four folds. This MVPA yielded a whole brain map of object image decoding performance, based on which the 200 most informative LOC voxels (in native volume space) in terms of image identity information were selected from the previously established LOC masks. This was done to ensure that the final masks contain voxels which best discriminate between the different object images. Freesurfer 6.0 (‘recon-all’; Dale et al., 1999) was utilized to extract V1 labels (left and right) per subject based on their anatomical image. Subsequently, the obtained labels were transformed back to native space using ‘mri_label2vol’ and combined into a bilateral V1 mask. The same searchlight approach mentioned above was used to constrain the anatomical V1 masks to the 200 most informative V1 voxels concerning object identity decoding. To verify that our results were not unique to the specific (but arbitrary) ROI size, we repeated all ROI analyses with ROI masks ranging from 50 to 300 voxels in steps of 50 voxels.

#### Software

FSL 5.0.9 (FMRIB Software Library; Oxford, UK; www.fmrib.ox.ac.uk/fsl; Smith et al., 2004) was utilized for preprocessing and analysis of fMRI data. Additionally, custom Matlab (The MathWorks, Inc., Natick, Massachusetts, United States) and Python (Python Software Foundation) scripts were used for additional analyses, data extraction, statistical tests, and plotting of results. The following toolboxes were used: NumPy (van der Walt et al., 2011), SciPy (Jones et al., 2001), Matplotlib (Hunter, 2007) and Scikit-learn (Pedregosa et al., 2011). Whole-brain results are displayed using Slice Display (Zandbelt, 2017) using a dual-coding data visualization approach (Allen et al., 2012), with color indicating the parameter estimates and opacity the associated z statistics. Additionally, MRIcroGL (www.mccauslandcenter.sc.edu/mricrogl) was used to 3D render the whole-brain results. Stimulus presentation was done using Presentation^®^ software (version 18.3, Neurobehavioral Systems, Inc., Berkeley, CA).

## Results

### Expectation suppression throughout the ventral visual stream

We first examined expectation suppression within our a priori defined ROIs, V1 and object-selective LOC. We observed a significantly larger BOLD response to unexpected compared to expected image pairs, both in V1 (*t*(21) = 3.20, *p* = 0.004, Cohen’s *d*_z_ = 0.68, Figure 2C) and object-selective LOC (*t*(21) = 5.03, *p* = 5.6e-5, Cohen’s *d*_z_ = 1.07, Figure 2C). To ensure that the results are not dependent on the (arbitrarily chosen) mask size of the ROIs, the analyses were repeated for ROIs of sizes between 50-300 voxels (691-4147mm^3^); the direction and statistical significance of all effects was identical for all ROI sizes.

**Figure 2.**
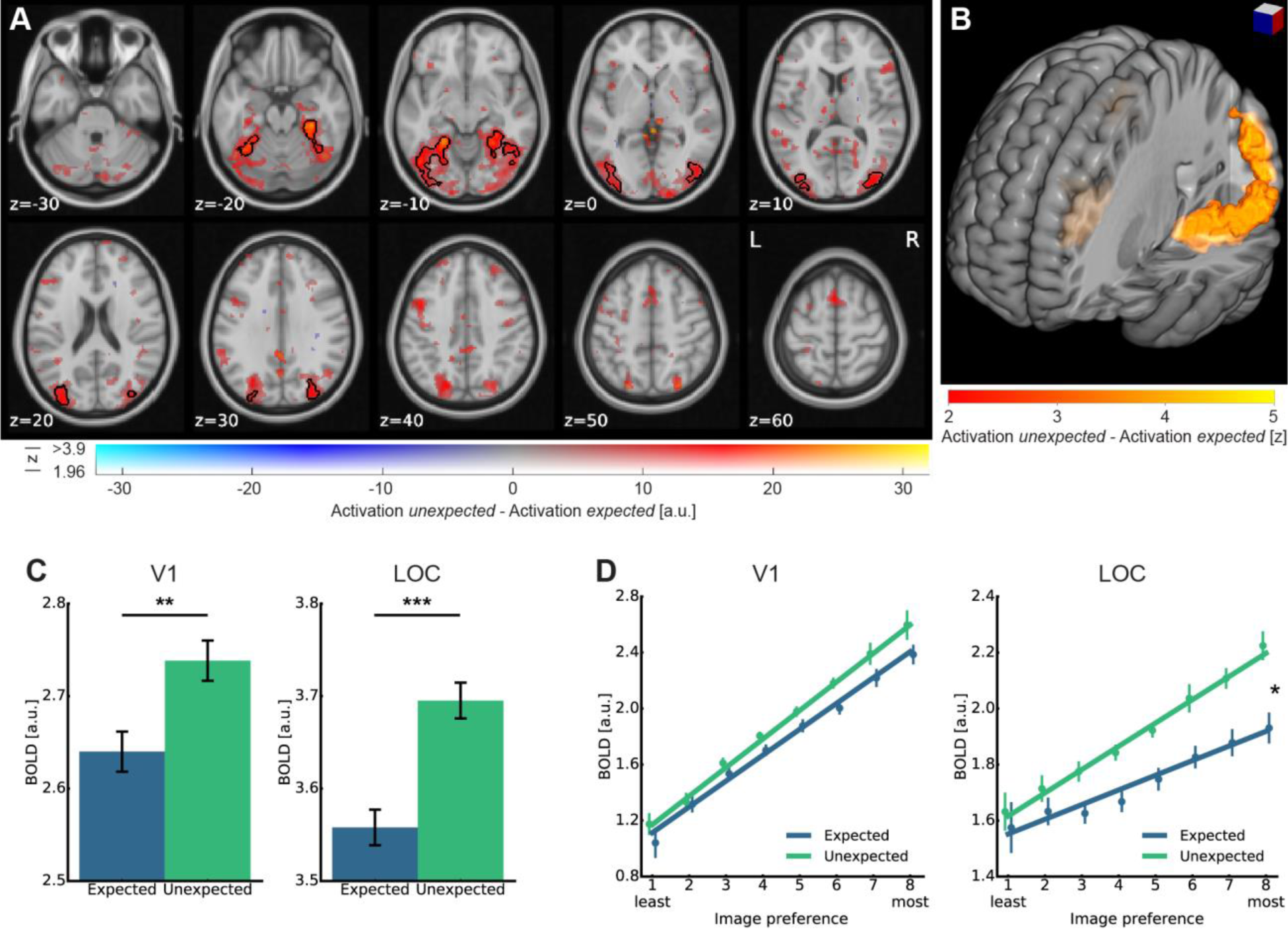
**(A)** Expectation suppression throughout the ventral visual stream. Displayed are parameter estimates for unexpected image pairs minus expected pairs overlaid on the MNI152 2mm template. Color represents the parameter estimates, with red-yellow clusters indicating expectation suppression, and opacity depicting the associated z statistics. Black contours outline statistically significant clusters (GRF cluster corrected), which include significant expectation suppression in superior and inferior divisions of LOC, temporal occipital fusiform cortex, and posterior parahippocampal gyrus. **(B)** Expectation suppression (cluster corrected) displayed over the 3D rendered MNI152 template brain. Color indicates z statistics of the expectation suppression contrast. Visible in the left hemisphere is one contiguous cluster, showing significant expectation suppression, including superior and inferior divisions of LOC, TOFC, and posterior PHG. The corresponding cluster in the right hemisphere (also see panel A) is occluded here. **(C)** Expectation suppression within V1 and object-selective LOC. Displayed are parameter estimates ± within-subject standard error for responses to expected and unexpected images pairs. In both ROIs, V1 (left bar plot) and LOC (right bar plot), BOLD responses to unexpected image pairs were significantly stronger than to expected image pairs. **(D)** Image preference analysis results in V1 and object-selective LOC. Parameter estimates ± within-subject standard error are displayed as a function of voxel-wise image preference, ranked from the least to the most preferred image rank based on the functional localizer. Superimposed is the mean regression line fit of the subject-wise regressions for expected and unexpected image pairs separately (see *Methods*). The left line plot shows responses to expected and unexpected image pairs within the V1 ROI. The fitted regression lines for expected and unexpected are parallel; i.e. no difference in slopes. The right plot displays image preference results for object-selective LOC, showing a steeper slope for the unexpected image pair regression line compared to the corresponding expected image pair regression line. * p <.05. ** p < .01, *** p < .001.

A whole-brain analysis, investigating effects of perceptual expectation across the brain, revealed an extended statistically significant cluster (Figure 2A, black contours) of expectation suppression across the ventral visual stream. Cortical areas showing significant expectation suppression included large parts of superior and inferior bilateral object-selective LOC, bilateral temporal occipital fusiform cortex (TOFC), and right posterior parahippocampal gyrus (PHG). In fact, the observed expectation suppression effect in the left hemisphere consisted of one large statistically significant cluster, extending throughout most of the ventral visual stream, as can be seen in Figure 2B. The corresponding significant effect in the right hemisphere, occluded in Figure 2B, consists of two large clusters.

Next, we assessed the neural effect of the conditional probability conditions within V1 and LOC. While this analysis confirmed a weaker response for expected items in V1 (*F*(1,21) = 6.39, *p* = 0.020) and LOC (*F*(1,21) = 19.50, *p* = 2.4e-4), there was no significant modulation by conditional probability, nor an interaction between conditional probability and expectation in either V1 (conditional probability: *F*(2,42) = 2.02, *p* = 0.145; interaction: *F*(2,42) = 1.19, *p* = 0.315) or LOC (conditional probability: *F*(2,42) = 1.90, *p* = 0.162; interaction: *F*(2,42) = 0.92, *p* = 0.407). We therefore collapse across the three different conditional probability conditions for all subsequent analyses.

### Perceptual expectations dampen sensory representation in LOC

To examine whether sharpening or dampening of sensory representations underlies expectation suppression in V1 and LOC, an image preference analysis was conducted. In short, BOLD responses were regressed on image preference rank, with dampening predicting a steeper slope for unexpected compared expected images and sharpening predicting the opposite (see *Methods* for details). Results, depicted in Figure 2D, reveal positive slopes within V1 (expected: *t*(21) = 9.11, *p* = 9.6e-9; V1 unexpected: *t*(21) = 9.90, *p* = 2.3e-9), as well as in LOC (expected: *t*(21) = 3.39, *p* = 0.003; LOC unexpected: *t*(21) = 7.14, *p* = 4.8e-7), confirming that the image preference ranking from the functional localizer data generalized to the main task. This indicates a stable, reproducible sensory code and allows for an analysis of the difference in slopes between expected and unexpected image pairs. Crucially, image preference slopes were significantly steeper for unexpected than expected image pairs in LOC (*t*(21) = 2.18, *p* = 0.041, Cohen’s **d*_z_=* 0.47), but not in V1 (*t*(21) = 1.20, *p* = 0.242). This means that the amount of expectation suppression (i.e. the difference in the two regression lines in Figure 2D) increased with the image preference rank in object-selective LOC, but not in V1. A control analysis confirmed that the results were independent of the number of voxels in the respective ROIs (mask sizes 50-300 voxels).

In a complementary analysis, we reasoned that if the reduced activity for expected items is associated with a reduction of noise (sharpening), it is expected to be associated with an increase in classification accuracy in a MVPA (Kok et al. 2012a). Conversely, a dampening of the representation is predicted to be associated with a decrease in classification accuracy for expected image pairs (Kumar et al., 2017). Generally, image identity could be classified well above chance (12.5%) in V1 (expected: 27.9%, *t*(21) = 10.89, *p* = 4.3e-10; unexpected: 30.2%, *t*(21) = 15.70, *p* = 4.5e-13), and LOC (expected: 18.5%, *t*(21) = 5.69, *p* = 1.2e-5; unexpected: 19.5%, *t*(21) = 6.76, *p* = 1.1e-6). While a trend towards better decoding performance for unexpected images was indeed visible in both ROIs, in line with dampening of the sensory response, this difference was not statistically significant (V1: *t*(21) = 1.93, *p* = 0.067; LOC: *t*(21) = 1.16, *p* =0.260).

### Expectation facilitates image categorization

In order to assess whether concurrent to the described neural effects also behavioral benefits of expectation are evident, data from the categorization task was analyzed. Results demonstrate that participants categorized expected trailing images faster (*M* = 524.4 ms, *SEM* = 3.8 ms) than unexpected items (*M* = 537.4 ms, *SEM* = 3.8; *t*(21) = 2.40, *p* = 0.026, Cohen’s *d*_z_ = 0.51; Figure 3A). A similar, albeit not statistically significant trend (*t*(21) = 1.19, *p* = 0.247) was visible in terms of error rates (Figure 3B). Analysis of the questionnaires showed that on average participants correctly identified 4.0 ± 2.3 (± SD) of the eight image pairs.

**Figure 3.**
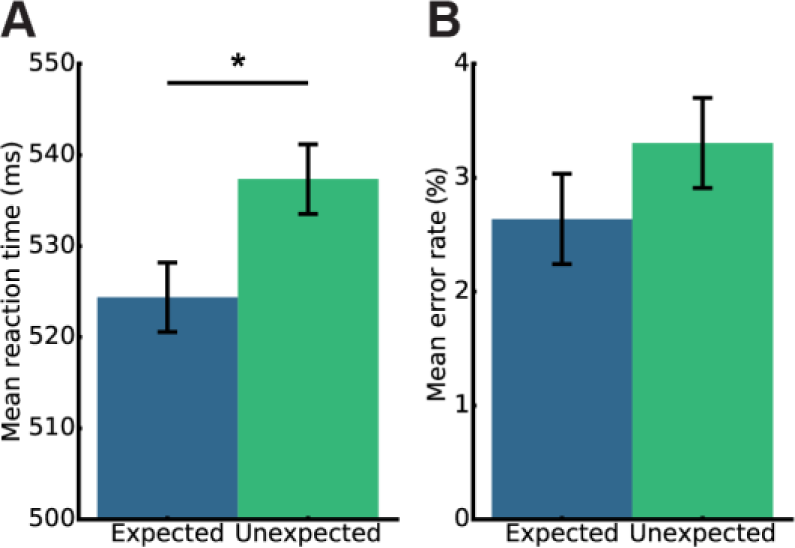
Behavioral data analysis froM the categorization task indicates incidental learning of image transitions. Mean values ± within-subject standard error are shown. (**A**) Shows mean RT to expected and unexpected trailing images. RTs were significantly faster to expected trailing images compared to unexpected images. (**B**) Shows the corresponding mean error rates. * p < .05.

### Spatial extent of expectation suppression in V1

In a post hoc analysis we investigated whether the expectation suppression effect in V1 was spatially unspecific, or constrained to regions activated by the object stimuli. The reasoning was that a spatially unspecific effect indicates that at least part of the observed expectation suppression may be due to arousal changes in response to unexpected compared to expected trailing images, while a constrained effect may point towards a spatially specific top-down modulation. To investigate this, the amount of expectation suppression was compared between voxels significantly activated by object stimuli and those that were not. The split into activated and not activated voxels was performed using data froM the functional localizer, with activated voxels being defined as all voxels within anatomically defined V1 which exhibited a significant activation by object images (z > 1.96; i.e. p < 0.05, two-sided), while non-activated voxels were defined as voxels displaying no significant activation, nor deactivation (-1.96 < z < 1.96). Both activated and non-activated voxels showed evidence of expectation suppression (activated voxels: *t*(21) = 3.01, *p* = 0.007, Cohen’s *d*_z_ = 0.64; non-activated voxels: *t*(21) = 2.17, *p* = 0.041, Cohen’s *d*_z_ = 0.46). While expectation suppression was numerically stronger in voxels that were activated by the stimuli than in non-activated voxels, this difference was not statistically significant (*t*(21) = 1.09, *p* = 0.286).

## Discussion

We set out to investigate the neural effects of perceptual expectation and demonstrated that, after incidental learning of transitional probabilities of object images, expectation suppression is evident throughout the human ventral visual stream. Importantly, the amount of expectation suppression scaled positively with image preference in LOC, suggesting that dampened sensory representations underlie expectation suppression in object-selective areas, in line with results from monkey IT (Meyer and Olson, 2011; Kumar et al., 2017). In contrast, results in V1 did not exhibit scaling or sharpening of representations, but rather stimulus unspecific expectation suppression.

### Dampening of sensory representation in object-selective cortex

The suppression of expected stimuli, evident throughout the ventral visual stream in the present study, extends and supports previous research showing expectation suppression in early visual areas (Kok et al., 2012a; St. John-Saaltink et al., 2015) and monkey IT (Meyer and Olson, 2011; Kaposvari et al., 2016). The observed suppression may constitute an efficient and adaptive processing strategy, which filters out predictable, irrelevant objects froM the environment. Conversely, the stronger response to unexpected objects may serve to render unexpected stimuli more salient. This surprise response to unexpected stimuli may draw attention towards these stimuli, as also reasoned by Meyer and Olson (2011). Such capture of attention is adaptive since unexpected events may provide particularly relevant information. It is important to note that the utilized paradigM did not manipulate attention towards expected or unexpected stimuli in a top-down fashion. In fact, unexpected and expected stimuli were only distinguishable by the context in which they occurred. Therefore, if unexpected stimuli do indeed automatically capture attention (Brockmole and Boot, 2009; Howard and Holcombe, 2010), then any attentional modulation must follow the expectation effect, and not vice versa.

Furthermore, we showed that the amount of expectation suppression scales with image preference in object-selective LOC, as also demonstrated in monkey IT (Meyer and Olson, 2011). Scaling indicates that expectation suppression in object-selective areas does *not* merely signal an unspecific surprise response, but rather that sensory representations are dampened by expectations, since the neural population most responsive to the expected stimulus is also most suppressed. A dampening of sensory representations is in line with an adaptive mechanism, which filters out behaviorally irrelevant, predictable objects froM the environment. Unlike in LOC, expectation suppression in V1 was stimulus unspecific; i.e. neither scaling, nor sharpening was observed. These results cannot be explained by the absence of image preference in V1 for the utilized stimuli, as the preference ranking itself was reliable. Since a stimulus unspecific suppression was evident in V1, it is possible that object specific expectations were resolved at a higher level in the cortical hierarchy and only the results of the prediction (expected or unexpected) was relayed to V1 as feedback. Given that expectation suppression was present in stimulus driven voxels, but to a lesser degree also in non-stimulus driven voxels, it seems plausible that expectation suppression in V1 arose as a combination of spatially unspecific arousal changes across V1 and stimulus unspecific, but spatially specific top-down modulations froM higher visual area, such as LOC.

If expectation suppression, and the underlying representational dampening, does in fact represent an adaptive neural strategy one may expect behavioral benefits to correlate with the neural effects. Although we observed behavioral benefits for expected stimuli during the categorization task, the present study cannot answer whether expectation suppression is associated with behavioral benefits, since during the fMRI task, and central to the interpretation above, expectations were task irrelevant. Task relevant predictions, necessary in order to investigate this question, may in turn change the underlying neural dynamics. In fact, it has been suggested that, at least in early visual areas, attention can reverse the suppressive effect of expectation (Kok et al., 2012b).

Finally, the present results appear to be at odds with a previous study that observed a sharpening of the sensory population response in V1 by expectation (Kok et al., 2012a). While there is a multitude of differences between the two studies, making it impossible to isolate one factor as definitive source of the discrepancy, we briefly discuss two aspects that may be particularly relevant to consider. Firstly, the two studies employed different stimuli (object vs. grating stimuli), tailored to investigate the population response in different areas of the visual hierarchy (LOC vs. V1). Given that we did not find a sharpening, nor dampening, of representations in V1 the opposite results cannot be explained by a general difference between the sensory areas, but rather an interaction between stimulus type and sensory area. Secondly, there are profound differences between the studies in task demands. In the current study, we examined neural activity elicited by expected and unexpected *non-target* stimuli, i.e. stimuli that did not require a response by the observer. On the other hand, all stimuli in Kok et al. (2012a) were *target* stimuli, requiring a discrimination judgment by the observers. Given that attentional selection is known to sharpen stimulus representations (Serences et al., 2009), this difference in task setup could be a relevant factor in explaining the opposite results between the studies.

### Prediction errors and predictive coding

Within a hierarchical predictive coding framework, prior expectations about an upcoming stimulus act as top-down signals predicting the bottom-up input based on generative models of the agent (Friston, 2005). These predictions are then compared to the actual bottom-up input resulting in a mismatch signal, the prediction error (PE). Expectation suppression, as evident in the present data, and previously suggested by others (e.g. Blank and Davis, 2016; den Ouden et al., 2012; Kok et al., 2012a), matches the properties of a PE signal. That is, the ensuing PE is smaller for expected compared to unexpected trailing images, since the mismatch between prediction and input is smaller, thus resulting in expectation suppression. We found such suppression throughout the ventral visual stream, within the same paradigm, corroborating the notion that perceptual PEs are a common mechanisM throughout sensory cortex (den Ouden et al., 2012). Crucially, the exact properties of PEs have been suggested to depend on the functional role of the cortical areas in which they arise (den Ouden et al., 2012), which the present data support by providing evidence for scaling in LOC and an unspecific suppression in V1. Furthermore, a dampening of object representations in LOC, and the associated trend towards superior decoding performances of unexpected images, can be explained within predictive coding as a result of the stronger and prolonged resolution of prediction errors elicited by unexpected images. Moreover, the observed scaling effect naturally follows froM expectation suppression conceptualized as PE, since a PE signal would be expected to scale with the strength of the prediction of a stimulus in the underlying neural population - i.e. the stronger a prediction within a neural population, the stronger the ensuing mismatch when the prediction is violated.

### No systematic modulation of expectation suppression by conditional probability

The present results do not provide evidence for a systematic modulation of expectation suppression by conditional probabilities. This is somewhat surprising given that a modulation has been demonstrated in monkey TE (Ramachandran et al., 2016). Furthermore, it is only by virtue of the difference in conditional probability that a trailing image can be considered expected or unexpected. Thus, by its nature expectation suppression should be sensitivity to conditional probability. We believe that this null result arises due to a lack of sensitivity of the associated analysis. The complexity of the transition matrix and the relatively small difference in conditional probability between the conditions, as well as the split of the available data into the three conditions may have all led to a reduction in sensitivity. Thus, to further elucidate the nature of expectation suppression future research in humans is required, possibly utilizing simplified paradigms or extended exposure to the image transitions.

## Conclusion

Taken together, our results demonstrate that expectation suppression is a wide-spread neural mechanisM of perceptual expectation, which scales with image preference in object-selective LOC, but not V1. Perceptual expectations thus lead to a selective dampening of sensory representations in object-selective cortex.

## Acknowledgements

We thank Alexis Pérez-Bellido and Micha Heilbron for helpful suggestions on the manuscript. This work was supported by The Netherlands Organisation for Scientific Research (NWO Vidi grant 452-13-016 awarded to FPdL) and the EC Horizon 2020 PrograM (ERC starting grant 678286, ‘Contextvision’ awarded to FPdL).

